# Mimicking of tau hyperphosphorylation in GABAergic motoneurons of *C. elegans* induces severe peripheral and neuronal alterations

**DOI:** 10.1101/2023.07.04.547705

**Authors:** Audrey Labarre, Émilien Schramm, Julie Pilliod, Samuel Boyer, Marianne Lapointe, Claudia Maios, Nicole Leclerc, J. Alex Parker

## Abstract

In several neurodegenerative diseases including Alzheimer’s disease (AD), tau, a microtubule-associated protein (MAP) enriched in the axon, becomes hyperphosphorylated, detaches from microtubules, redistributes to the somato-dendritic compartment and self-aggregates. The mechanisms leading to neuronal dysfunction and death by tau pathology remain to be fully elucidated. *C. elegans* has been successfully used by several groups including ours to identify mechanisms involved in neurodegeneration. We generated three strains, one overexpressing wild-type human tau (WT Tau), one a tau mutant mimicking hyperphosphorylation (hyperP Tau) and one preventing phosphorylation (hypoP Tau) in GABA motor neurons. A significant reduction of body size and egg laying was noted in these tau strains. Starting at day 1, we found that the worms overexpressing hyperP Tau were smaller than the N2 control strain and the worms either overexpressing WT Tau or hypoP Tau. Starting at day 5, the worms overexpressing WT Tau were smaller than control and the worms overexpressing hypoP Tau. Egg laying was reduced in both hyperP Tau and WT Tau worms. Survival was only decreased in WT Tau worms. Motility deficits were also observed. For age-dependent paralysis, a difference was noted between control and hyperP Tau. Swimming activity and speed were increased in hypoP Tau and decreased in hyperP Tau strains. Axonal integrity was altered in all tau strains. In the case of synaptic activity, at day 1, it was increased in the hypoP Tau strain and decreased in the hyperP Tau one. Collectively, our data revealed that overexpression of tau exerted neuronal and peripheral defects indicating that tau dysfunction could affect cell-cell communication.

## Introduction

Tau is an axonal MAP that becomes hyperphosphorylated, accumulates in the somato-dendritic compartment and self-aggregates into insoluble filaments called paired helical filaments (PHFs) forming the neurofibrillary tangles (NFTs) in AD (Ludin and Matus, 1993; Mandell and Banker, 1996; Lee et al., 2001; Cairns et al., 2007; Iqbal et al., 2016). Tau pathology is correlated to cognitive deficits in patients as demonstrated by histopathological examination of post-mortem brain and tau PET imaging (Tomlinson et al., 1970; Alafuzoff et al., 1987; Braak and Braak, 1991; Arriagada et al., 1992; Bierer et al., 1995; Ossenkoppele et al., 2016; Pontecorvo et al., 2019). No mutations in the tau gene have been found in AD patients but tau gene polymorphisms may be risk factors for sporadic AD (Schraen-Maschke et al., 2004). A duplication of the tau gene was recently correlated to an early-onset dementia with an AD clinical phenotype (Le Guennec et al., 2017). The contribution of tau dysfunction to neurodegeneration is further supported by the enrichment of tau genetic variants in patients suffering from frontotemporal lobar degeneration (FTLD-tau) (Cairns et al., 2007). Although all the above observations indicate that tau pathology is correlated to neurodegeneration, its precise role in this process remains elusive.

Several animal models have been produced to investigate the contribution of tau pathology to neurodegeneration, but no model has allowed complete elucidation of the mechanisms underlying tau toxicity. In most models, tau hyperphosphorylation is correlated to tau pathology and toxicity (Sanchez-Varo et al., 2022). It is believed that hyperphosphorylation contributes to tau detachment from microtubules, which is linked its aggregation (Iqbal et al., 2016). However, it is still unclear how tau hyperphosphorylation contributes to neuronal dysfunction. In pathological conditions and aging, some studies reported that tau hypophosphorylation preceded tau hyperphosphorylation (Wander et al., 2020). Our previous study has shown that neurons have mechanisms to maintain the constant global phosphorylation state of tau (Bertrand et al., 2010). One can speculate that these mechanisms are perturbed in pathological conditions leading to hypophosphorylation followed by hyperphosphorylation.

Based on the above observations, we developed *C. elegans* models either overexpressing a human tau mutant mimicking hyperphosphorylation (hyperP Tau), a non-phosphorylable mutant (hypoP Tau) or wild-type tau (WT Tau). Several models of tau transgenic *C. elegans* already exist (Pir et al., 2017). Pan-neuronal expression of human tau was associated with a progressive uncoordinated phenotype indicating neuronal defects. Accumulation of insoluble tau and neurodegeneration were noted with increasing age. More severe alterations were observed with tau mutants linked to fronto-temporal lobar degeneration (FTLD-tau) (Pir et al., 2017; Aquino Nunez et al., 2022). In previous studies where the expression of human tau was pan-neuronal, worms were often very sick from an early age compromising the investigation of the mechanisms involved in neuronal dysfunction induced by its pathology. Because of this, in our models, human tau was expressed only in the 26 GABAergic motoneurons. The neurodegeneration of these neurons induces a paralysis phenotype that may be useful for genetic and drug screening applications. We successfully used such an approach to investigate the mechanisms involved in other neurodegenerative conditions such Huntington’s disease and amyotrophic lateral sclerosis (Aggad et al., 2014; Labarre et al., 2021; Labarre et al., 2022). In a previous study, human tau mutant mimicking phosphorylation, and one preventing it were pan-neuronally expressed in *C. elegans* (Brandt et al. 2009). WT Tau and the mutant mimicking phosphorylation presented an uncoordinated locomotion phenotype correlated with the presence of axonal gaps. A recent study performed on tau isolated from AD brain reported that some phosphorylation sites are linked to tau seeding competency meaning that their phosphorylation increases tau aggregation (Wesseling et al., 2020). In the Brandt’s study, only 4 of the 9 phosphorylation sites mimicking phosphorylation were seeding competent. In our models, we mutated 12 sites, 8 seeding competent and 4 seeding incompetent. The 4 sites seeding incompetent are detected at a pre-tangle stage in AD brain. We found that the worms overexpressing tau mutant mimicking hyperphophorylation (hyperP Tau) were smaller than the controls and the worms either overexpressing the tau mutant preventing phosphorylation (hypoP Tau) and WT Tau. For both hyperP Tau and WT Tau worms, egg laying was reduced. Motility deficits were observed in hyperP Tau and hypoP Tau worms. For age-dependent paralysis, a difference was noted between the control and hyperP Tau strains. Swimming activity and speed were increased in hypoP Tau and decreased in hyperP Tau strains. Axonal integrity was altered in all tau strains. In the case of synaptic activity, at day 1, it was increased in the hypoP Tau strain and decreased in the hyperP Tau one.

## Results

We generated three tau *C. elegans* strains, one overexpressing the 0N4R isoform of human tau (WT Tau), one 0N4R mutant containing 12 phosphorylation sites that were mutated to alanine to prevent phosphorylation (hypoP Tau) and one 0N4R mutant containing the same 12 phosphorylation sites mutated in glutamate to mimic phosphorylation (hyperP Tau) (Figure 1A and S2). The protein levels of human tau were analyzed by western blotting (Figure 1B). A band corresponding to human tau was only found in strains overexpressing either WT Tau or a tau mutant but not in the N2 control strain. A band of high molecular weight was consistently observed in the hyper-tau strain indicating the presence of SDS-resistant tau oligomers. Such a band was occasionally observed in the WT Tau and hypoP Tau strains.

**Figure 1.**
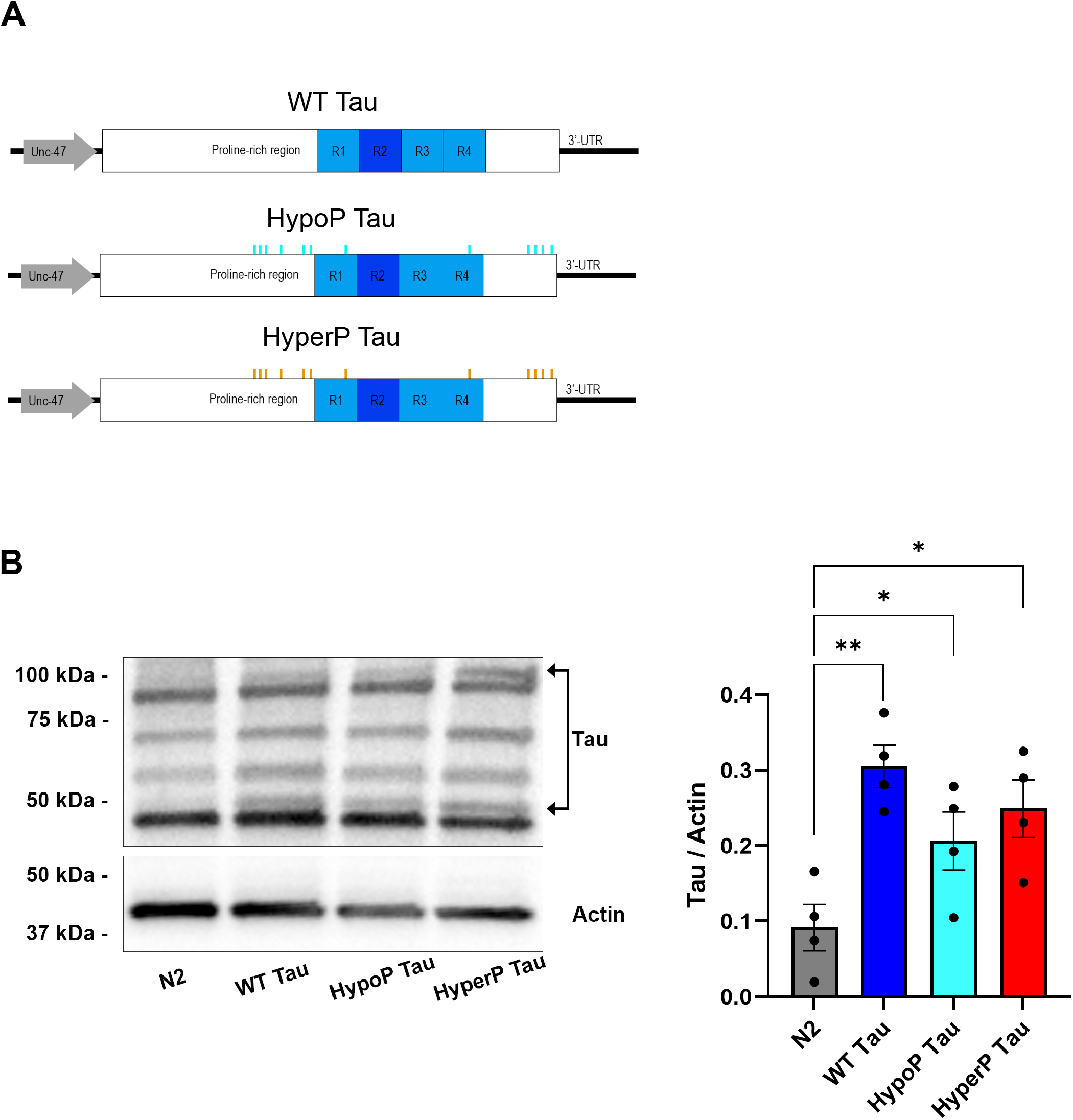
Tau transgene constructs and protein levels in *C. elegans*. **(A)** Schematic representation of constructs containing human Tau under the *unc-47* promotor. The following full-length Tau constructs were cloned for expression in the GABAergic motor neurons and injected into *C. elegans*: wild type (WT) Tau, hypophosphorylated (HypoP) Tau containing 12 mutated sites to alanine, and hyperphosphorylated (HyperP) Tau containing 12 mutated sites to glutamate. **(B)** Representative Western blot of total proteins from non-transgenic N2, WT Tau, HypoP Tau and HyperP Tau worms. Total Tau was normalized to actin, the loading reference. Densitometry quantification of Tau/Actin ratio showed that Tau protein is expressed in all transgenics. One-way ANOVA was performed with N=4. Data represent mean±SEM. *p < 0.05; **p < 0.01. R1: microtubule-binding repeats region 1; R2: microtubule-binding repeats region 2; R3: microtubule-binding repeats region 3; R4: microtubule-binding repeats region 4.

Representative images of tau strains revealed that hyperP Tau worms were smaller than N2, WT Tau and hypoP Tau animals at day 1 (Figure 2 A and B). At day 5, although hyperP Tau worms remained the shortest, WT Tau worms were also shorter than the N2 strain (Figure 2C). At day 9, the length of WT Tau worms continued to decrease and became not statistically different from that of hyperP Tau (Figure 2D). To eliminate the possibility that hyperP Tau and WT Tau worms were shorter because their feeding capacity was impaired, we examined their pharyngeal pumping activity (Figure S3). The mutant *eat-2* presenting very low pumping activity was using as a negative control, and we observed that pumping activity was normal for all tau strains.

**Figure 2.**
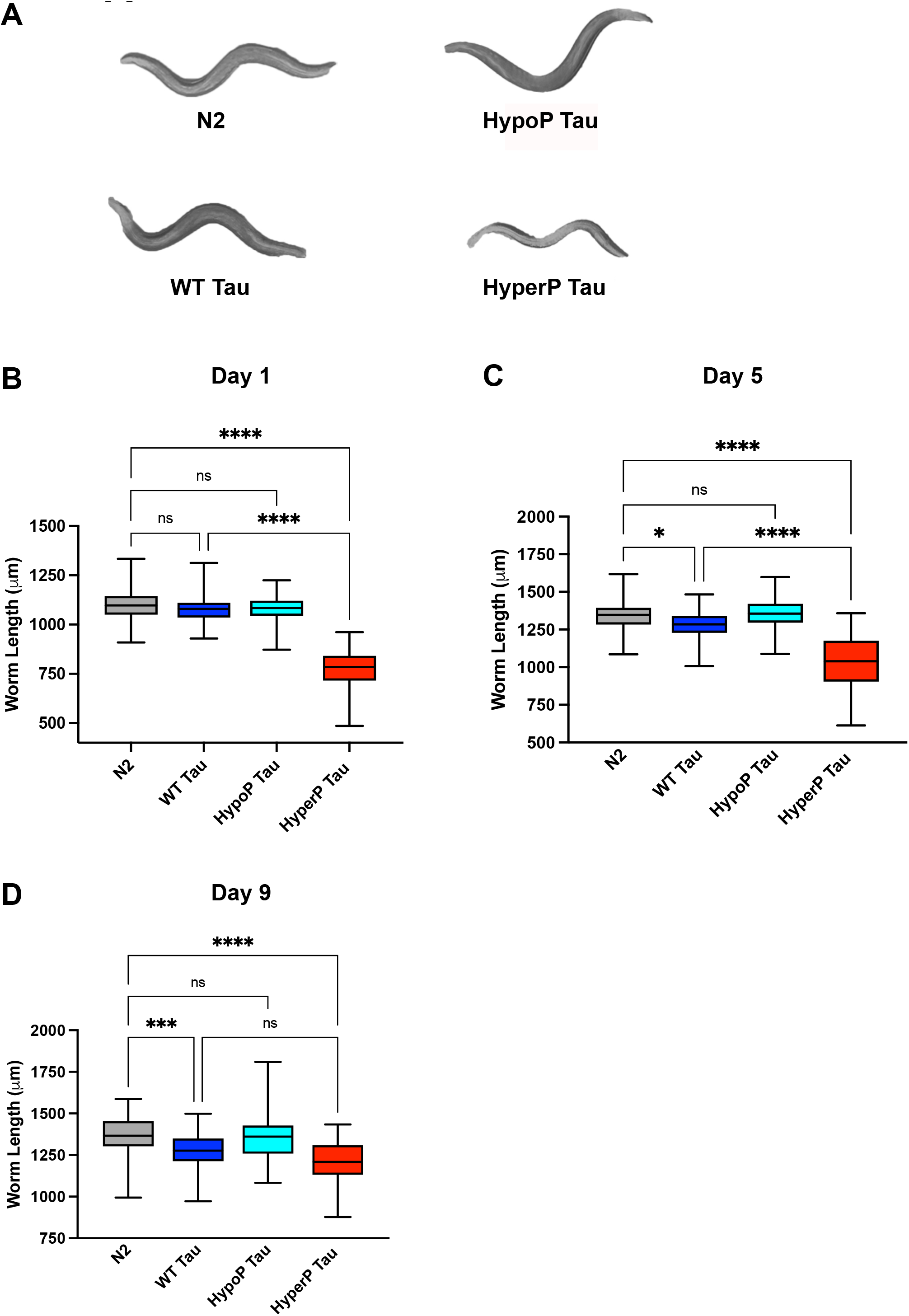
GABAergic HyperP Tau expression induces decreased body size in *C. elegans*. **(A)** Representative images of N2, WT Tau, HypoP Tau and HyperP Tau worms at day 1 of adulthood. Quantification of body length at day 1 **(B)**, day 5 **(C)** and day 9 **(D)** for all tau strains. HyperP Tau worms show decreased body size compared to N2 and other Tau transgenics. One-way ANOVA was performed. Between 42 and 82 individual worms were tested for each condition. For boxplots, minimum, first quartile, median, third quartile, and maximum are shown. *p < 0.05; ***p < 0.001; ****p < 0.0001.

The total progeny of tau strains was also examined (Figure 3A and B). Total progeny was decreased for both WT Tau and hyperP Tau compared to N2 although their profiles of egg laying were different. For the first two days, egg laying was lower for hyperP Tau than WT Tau. In the following days, hyperP Tau presented a progressive decrease of egg laying. In the case of WT Tau, egg laying stopped at day 4. Next we investigated the lifespan of tau strains but only the WT Tau strain showed a decrease of lifespan that was statistically different of the N2 strain (Figure 3C).

**Figure 3.**
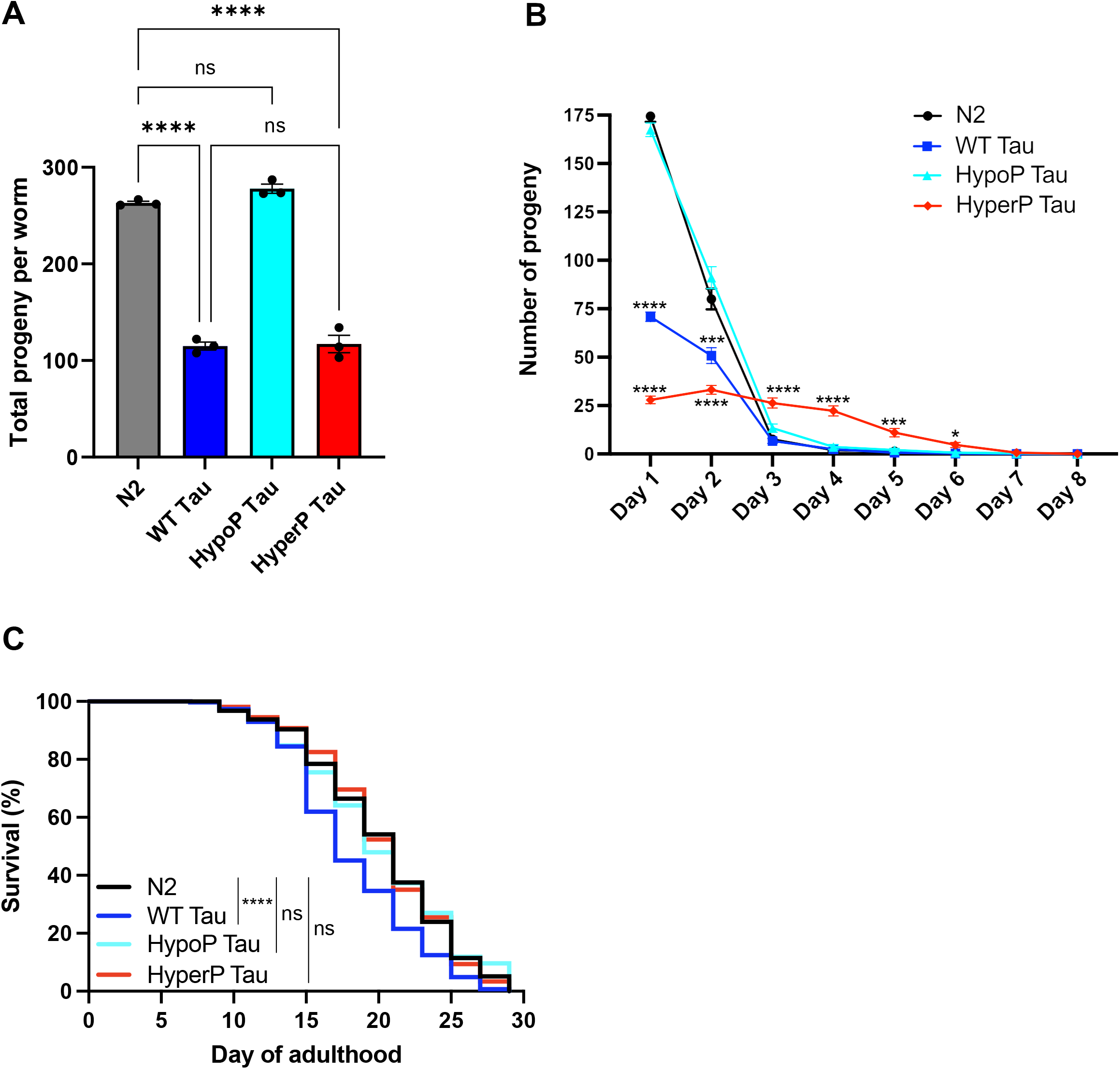
Tau expression decreases progeny and affects lifespan. **(A)** Quantification of total progeny per worm for N2, WT Tau, HypoP Tau and HyperP Tau animals. One-way ANOVA was performed. Data represent mean±SEM of N=3 (*n* between 27 and 30 worms total for each condition). **(B)** Quantification of daily progeny production per worm. Mixed-model ANOVA was performed. Data represent mean±SEM of N=3 (*n* between 27 and 30 worms total for each condition). Both WT Tau and HyperP Tau show decreased total progeny and decreased daily progeny production. **(C)** Worms were monitored from the adult stage and scored every two days for mortality. WT Tau worms have a shorter lifespan than N2 strain. Animals expressing HypoP and HyperP Tau transgenes have similar lifespan than N2. N=3 (*n*=270 worms total for each condition). Lifespan curve was generated and compared using the log-rank (Mantel–Cox) test. *p < 0.05; ***p < 0.001; ****p < 0.0001.

We then examined whether the significant peripheral defects noted in WT Tau and hyperP Tau strains were correlated to motility defects. Only hyperP Tau showed motility problems leading to age-dependent paralysis over 14 days (Figure 4A). To further investigate the motility of tau strains, swimming activity and speed were recorded using Wormtracker and Wormlab methods respectively (Schmeisser et al., 2017). At day1, an increase of activity was noted for hypoP Tau and a decrease for hyperP Tau compared to N2 (Figure 4B). No difference was noted for WT Tau. In the case of swimming speed, at day 1, an increase of speed was noted for hypoP Tau and a decrease for hyperP Tau (Figure 5A). This correlated with the increase of swimming activity for hypoP Tau and the decrease for hyperP Tau showed in Figure 4B. At day-5, we noted an increase of speed for WT Tau (Figure 5B). In the case of tau mutants, no significant difference was noted between hypoP Tau and N2, but hyperP Tau still presented a decreased speed compared to N2. At day 9, only hyperP Tau presented a difference with N2 (Figure 5C).

**Figure 4.**
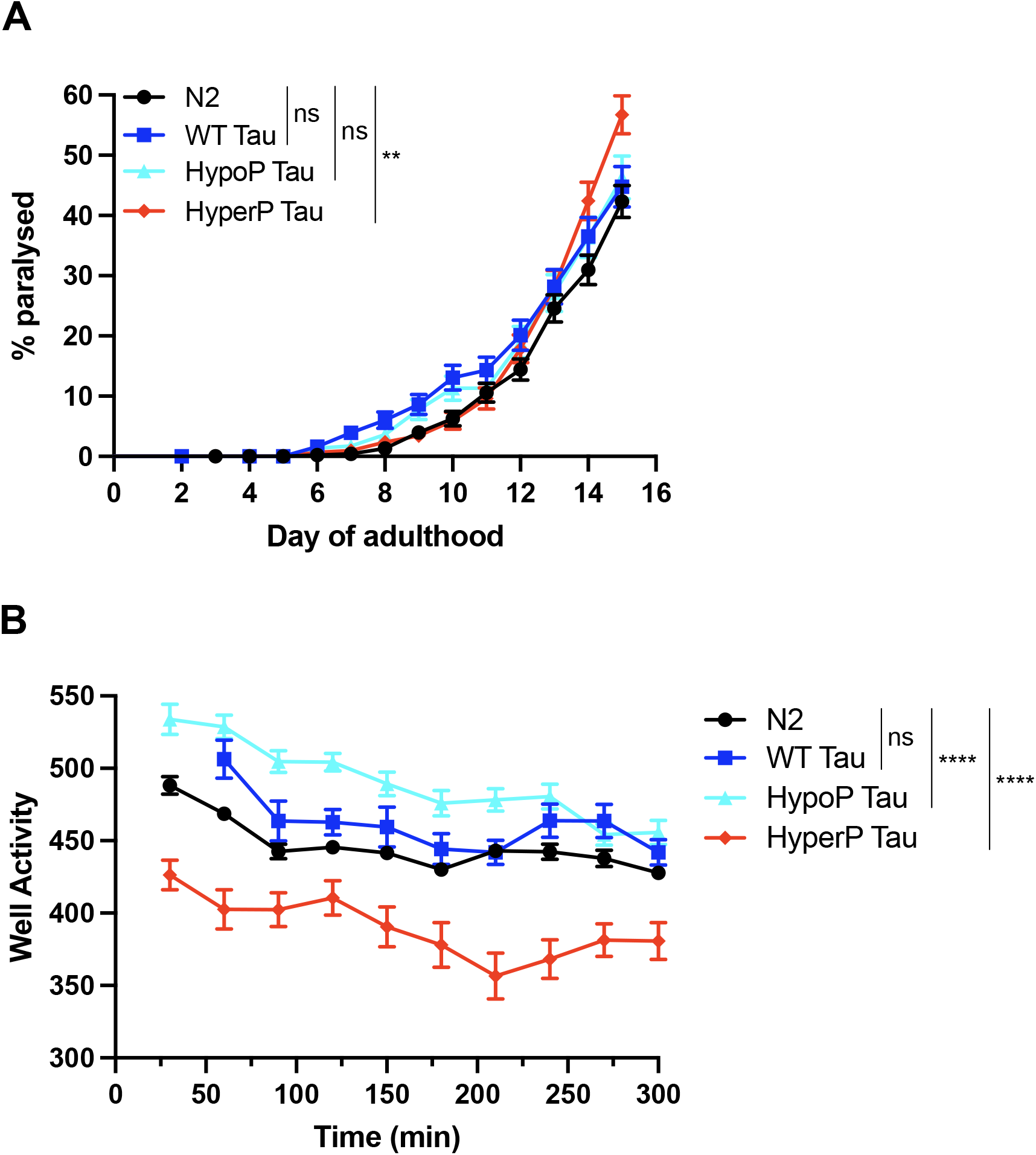
HyperP Tau causes age-dependent paralysis and motility impairment. **(A)** Worms were scored daily on solid media for paralysis phenotype, from day 1 to day 14. HyperP Tau transgenics show increased progressive paralysis over 14 days compared to N2, WT and HypoP Tau. N=3 (Between 315 and 502 individual worms were tested for each condition.). Paralysis curve was generated and compared using the log-rank (Mantel–Cox) test. **(B)** Motility of day 1 adult worms was assessed in liquid media, using a PhylumTech WMicrotracker One apparatus. HyperP Tau worms have decreased motility over 10 hours, while HypoP Tau have increased motility capacity. No significant difference was observed between WT Tau and N2 worms. Two-way ANOVA with a Tukey’s multiple comparisons test was performed. N=6 (*n*=540 worms total for each condition). **p < 0.01; ****p < 0.0001.

**Figure 5.**
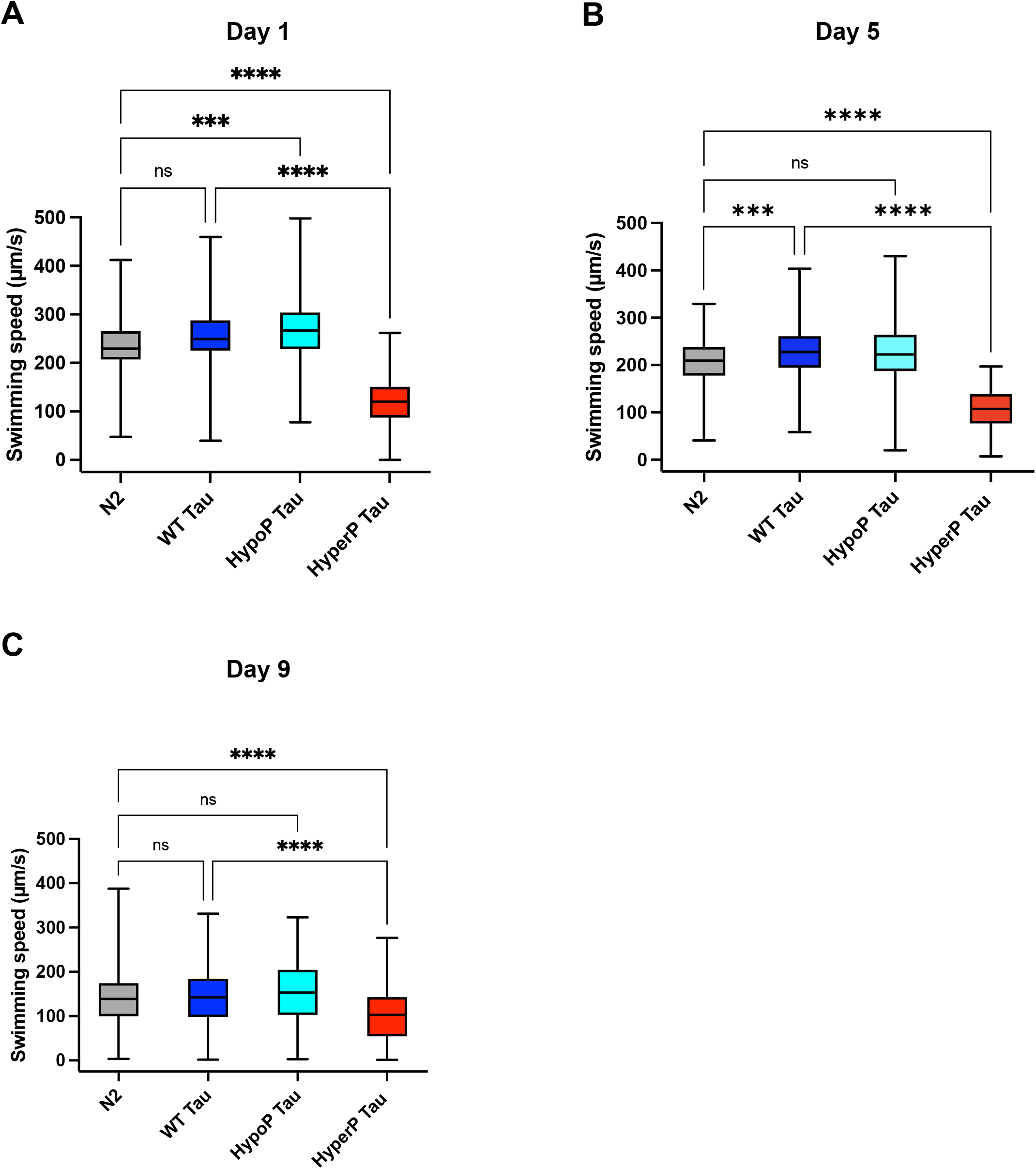
HyperP Tau transgenic worms have decreased swimming speed. Quantification of swimming speed at day 1 **(A)**, day 5 **(B)** and day 9 **(C)** for all the Tau strains. HyperP Tau transgenics show decreased swimming speed when compared to N2 and other Tau transgenics. HypoP Tau worms showed increased swimming speed at day 1 of adulthood. One-way ANOVA was performed. Between 88 and 187 individual worms were tested for each condition. For boxplots, minimum, first quartile, median, third quartile, and maximum are shown. ***p < 0.001; ****p < 0.0001.

We also measured the wave initiation rate (number of waves/minute) (Figure S4). At day 1, hyperP Tau presented the lowest rate compared to N2 (Figure S4A). In the case of hypoP Tau, an increase was noted. At day 5, only hyperP Tau presented a significant decrease compared to WT Tau and N2 (Figure S4B). At day 9, hypeP Tau still showed a decrease compared to N2 (Figure S4C). From the above observations, we could conclude that the motility and the peripheral defects were not linked since hypoP Tau did present motility defects but not the peripheral ones.

The paralysis and swimming defects noted for hyperP Tau could indicate that motor neurons were affected. This is consistent with previous studies reporting that pan-neuronal overexpression of human tau induced neurodegeneration and/or defects in axonal outgrowth (Brandt et al., 2009). The axonal integrity of our tau strains was examined (Figure 6). As early as day 1, axonal gaps were noted in all our tau strains (Figure 6A and B). Similar observations were made at days 5 and 9 (Figure 6C and D). At day 9, hyperP Tau presented a significant increase of axonal gaps compared to WT Tau and hypoP Tau (Figure 6D). Interestingly, at day 9, only hyperP Tau presented a decrease of swimming speed that was statistically different from that of the N2 strain.

**Figure 6.**
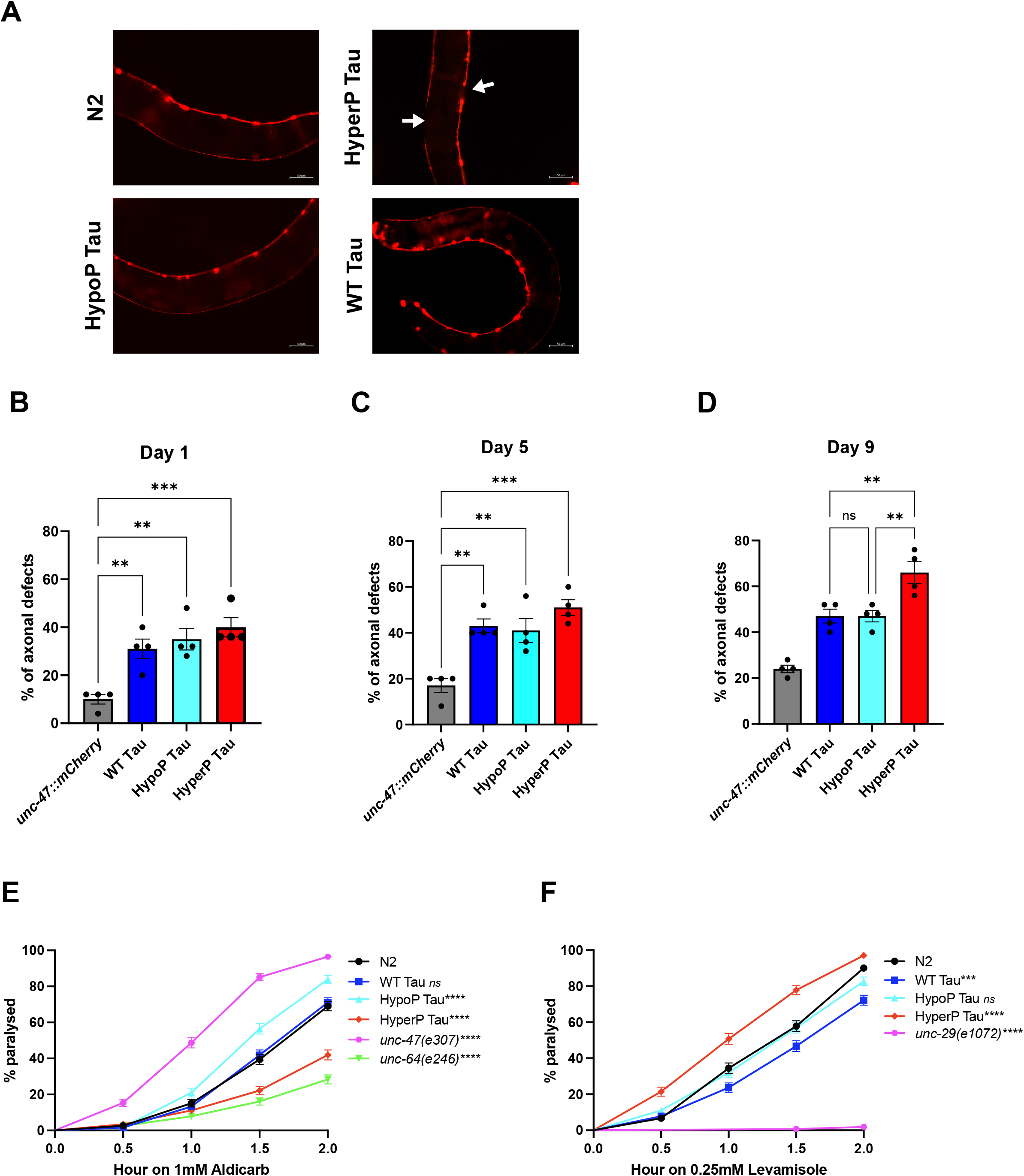
Neuronal tau expression induces axonal defects and altered synaptic transmission. **(A)** Representative images of worms expressing mCherry in the GABAergic motor neurons. Gaps along neuronal processes are indicated by the arrows. Scale bar = 50 μm. All transgenics expressing Tau (WT, HypoP and HyperP) showed axonal defects at day 1 **(B)** and day 5 **(C)** of adulthood **(D)**. HyperP Tau worms showed increased gaps along processes when compared to the other Tau strains at day 9. **(E)** Synaptic cholinergic transmission was evaluated by exposing adult worms to the cholinesterase inhibitor aldicarb at day 1. HypoP Tau worms are hypersensitive to aldicarb while HyperP Tau animals are resistant to the treatment. **(F)** Expression of HyperP Tau in GABAergic neurons causes sensitivity to levamisole induced paralysis. For axonal defects assessment, 100 worms were tested for each condition. One-way ANOVA was performed. For aldicarb and levamisole assays, between 270 and 317 individual worms were tested for each condition. Both curves were generated and compared using the log-rank (Mantel–Cox) test. **p < 0.01; ***p < 0.001; ****p < 0.0001.

Collectively, our observations suggested that motility defects could be caused by the axonal alterations, which could result in synaptic dysfunction. Two drugs, aldicarb and levamisole, were used to examine synaptic activity. At day-1, aldicarb leading to the accumulation of acetylcholine at the synaptic cleft was used to determine whether tau strains presented synaptic alterations (Figure 6E), The mutant *unc-47*, hypersensitive to aldicarb was used as a positive control and the mutant *unc-64* resistant to aldicarb was used as a negative control. The sensitivity of WT Tau to aldicarb was similar to that of the N2 strain. Interestingly, hypoP Tau was the most sensitive to aldicarb whereas hyperP Tau showed lesser sensitivity. Taken together, the above observations indicated that both tau mutants had synaptic alterations, hypoP Tau presenting an increase of synaptic activity and hyperP Tau, a decrease. To determine whether, the synaptic alterations were induced by pre-synaptic deficits, levamisole, an agonist of acetylcholine receptors, was used (Figure 6F). The mutant *unc-29* was used as a control resistant to levamisole. No difference was noted between hypoP Tau and N2. HyperP Tau was more sensitive to levamisole than N2, hypoP Tau and WT Tau indicating that the synaptic alterations observed in this mutant were induced by pre-synaptic defects. The above observations indicated that hypoP Tau and hyperP Tau strains developed different synaptic alterations.

## Discussion

In the present study, we generated three novel tau strains to investigate the contribution of tau hyperphosphorylation to neurodegeneration. Significant peripheral and neuronal defects were noted with the tau mutant mimicking hyperphosphorylation (hypeP Tau). The peripheral defects were a decrease of body size and egg laying. A decrease of survival was only observed for the WT Tau strain whereas an age-dependent paralysis was only noted for the hyperP Tau strain. In the case of motility, at day 1, swimming activity and speed were increased in hypoP Tau and decreased in hypeP Tau strains. Synaptic activity was also affected in tau mutant strains. At day 1, it was increased in the hypoP Tau strain and decreased in the hyperP Tau one. Axonal integrity was altered in all tau strains. Collectively, our data revealed that depending on its phosphorylation state, human tau exerts distinct peripheral and neuronal defects in *C. elegans*.

HyperP Tau worms were the smallest starting at day 1 whereas WT Tau worms became shorter than controls at day 5. The fact that the worms overexpressing hypoP Tau did not show any body size alterations indicate that tau phosphorylation contributed to this phenotype. A reduction of body size was not reported in previously published tau *C. elegans* strains, in which tau overexpression was pan-neuronal (Brandt et al., 2009). One explanation could be that different populations of neurons exert opposite effects on body size, some increasing it and others decreasing it. Such a scenario was reported for longevity. Certain gustatory neurons inhibit longevity whereas others promote it (Alcedo and Kenyon, 2004). Our results indicate that GABAergic motoneurons would play an important role in full body size development. This role could involve the secretion of morphogens such as DBL-1 playing a key role in body size (Maduzia et al., 2005). It is important to mention that the expression of other proteins linked to neurodegenerative diseases such as TDP-43 and FUS in the GABA motor neurons did not alter body size indicating that this effect is specific to tau (Aggad et al., 2014; Labarre et al., 2021). We previously reported that hyperphosphorylated tau can attach to membranous organelles involved in secretion such as Golgi membranes and induce their fragmentation (Liazoghli et al., 2005; Farah et al., 2006). One can speculate that hyperP Tau and to lesser extent WT Tau could attach to these membranes and thereby perturb the secretion of morphogens involved in body size. The contribution of tau to axonal transport and synaptic function could also significantly perturb the secretion of these neurons (Kent et al., 2020). We and others have shown that tau can be secreted (Pernegre et al., 2019). Another possibility is that extracellular tau could interfere with the signaling of morphogens involved in body size.

Egg laying was reduced in both WT Tau and hyperP Tau worms. GABA was shown to regulate egg laying (da Silva et al., 2022). HyperP Tau and WT Tau could affect the release of GABA. Indeed, hyperphosphorylated Tau was shown in mice to impair synaptic vesicle fusion with the plasma membrane (McInnes et al., 2018). However, only hyperP Tau presented pre-synaptic defects in our strains. As mentioned above, another possibility is that the secretion of factors involved in egg laying could be altered in GABA neurons by hyperP Tau and WT Tau. No problems of body size and egg laying was observed in hypoP Tau strain indicating that tau hyperphosphorylation was directly involved in these alterations. Both body size and egg laying alterations indicate that tau could affect cell-cell communication, but the mechanisms remain to be elucidated.

The lifespan of the strain overexpressing WT Tau was reduced compared to the N2 one. A previous study reported that microtubule stability was involved in lifespan. The deletion of PTL-1, the *C. elegans* ortholog of tau was shown to reduce lifespan whereas the deletion of EFA-6, a negative regulator of microtubule growth, increased it (Chew et al., 2013; Xu et al., 2019). Interestingly, the deletion of PTL-1 microtubule-binding domain was sufficient to reduce lifespan (Chew et al., 2013). In the PTL-1 mutant, microtubules were longer as well as the cargo run length indicating that PTL-1 regulates microtubule organization as well as axonal transport (Yogev et al., 2016). PTL-1 and tau share a high sequence homology in their microtubule-binding domain (McDermott et al., 1996). It is possible that human tau impairs the microtubule binding of PTL-1 by sequestering it, which would result in a reduced lifespan. Indeed, in mice, human pathological hyperphosphorylated tau was shown to sequester endogenous murine tau (Xia et al., 2022). In a previous study where the expression of human tau was pan-neuronal in *C. elegans*, human wt-tau was reported to be hyperphosphorylated (Brandt et al., 2009). In the present study, because human tau was only expressed in GABAergic neurons, its protein levels were too low to examine its phosphorylation. There is no obvious reason to be believe that wt-tau does not undergo phosphorylation in our strains.. HypoP Tau was expected to present an increase binding to microtubules, and thereby increased microtubule stability as reported in Drosophila, which could have resulted in an increase of lifespan (Talmat-Amar et al., 2011). However, the lifespan of the hypoP Tau strain was similar to that of the N2 strain. From the above observations, one can speculate that the increase of lifespan observed in WT Tau strain might involve a mechanism other than microtubule stability. For example, the four cephalic astrocyte-like sheath glia cells were shown to regulate lifespan through the secretion of neuropeptides activating a cell nonautonomous ER stress response in peripheral tissues in *C. elegans* (Frakes et al., 2020). Similarly, neurons were shown to be involved in longevity by release of small clear synaptic vesicles containing neurotransmitters (Taylor and Dillin, 2013). In WT Tau strain, these signaling pathways might be altered because of a defect in secretion. As mentioned above, tau can bind to membranous organelles involved in secretion, which could perturb the process of secretion. These secretory pathways would not be altered in hyperP and hypoP Tau strains that did not show any lifespan reduction.

GABA motor neurons play an important role in locomotion. The hyperP Tau strain presented the strongest locomotion defects. Interestingly, all the strains presented axonal alterations indicating that these two events are not linked. In the case of swimming speed and activity, hypoP Tau presented an increase whereas it was the opposite for hyperP Tau indicating that hypoP Tau and hyperP Tau could affect distinctly synaptic functions as revealed by the aldicarb treatment. Interestingly, the GABA-ion channel mutant, *unc-49,* is more sensitive to levamisole like hyperP Tau (Richmond and Jorgensen, 1999; Vashlishan et al., 2008). This could indicate that hyperP Tau affects this channel. The synaptic alterations induced by human tau in *C. elegans* are not surprising since its worm ortholog, PTL-1, was also showed to be involved in synaptic function. The *ptl-1(ok621)* mutant was reported to be less sensitive to levamisole indicating that tau contributes to the levamisole sensitivity (Chew et al., 2013). For both WT Tau and hypoP Tau, their swimming speed varied between day 1 and 9, which could indicate that their effects on synaptic function were temporally modulated.

In all our strains, axonal gaps were noted as early as L4 stage indicating that human tau expression induced developmental defects. In a previous study, such defects were linked to the aggregation of hyperphosphorylated Tau (Brandt et al., 2009). In our strains at day 1, HyperP Tau appeared to be able to form tau oligomers as revealed by western blotting. However, we could not confirm its aggregation as previously reported because of its selective expression in GABAergic motoneurons rendering the detection of aggregates very difficult. In a previous study, axonal gaps were only noted with WT Tau and hyper-tau and not with hypoP Tau although its overexpression was the highest (Brandt et al., 2009). Such a discrepancy could be explained by the difference of hypoP Tau protein levels. Indeed, it was previously demonstrated that a tight regulation of PTL-1 protein levels was determinant in its role in lifespan (Chew et al., 2013). The fact that all tau strains presented axonal gaps indicate that tau aggregation might not be the sole reason. Another explanation could be that microtubule dynamics were perturbed in all tau strains compromising protein transport in the axon. Our results revealed that synaptic activity is altered in hypoP Tau and hyperP Tau strains. Based on this, synaptic dysfunction could be involved in the axonal defects observed in these strains. From the above observations, one can speculate that distinct mechanisms could account for the axonal defects in our strains. In hypoP Tau worms, no alteration of egg laying and body size was observed whereas axonal and synaptic defects were present. This indicates that different pathways are involved in the peripheral and neuronal alterations and that these pathways are differently affected by hyperP and hypoP Tau.

Our work in primary hippocampal neurons has demonstrated that the state of tau phosphorylation results from priming and feedback events (Bertrand et al., 2010). Consistent with this, master phosphorylation sites that determine the propagation of phosphorylation at several sites were recently identified in mouse brain (Stefanoska et al., 2022). Together these studies demonstrate that in normal conditions, neurons use different mechanisms to maintain the physiological phosphorylation state of tau. In pathological conditions, such mechanisms would be inefficient in re-establishing this state leading either to hypophosphorylation or hyperphosphorylation. In previous studies, hypophosphorylation of tau was reported to precede tau hyperphosphorylation in AD (Wander et al., 2020). Based on this, our models could be used to elucidate the mechanisms leading to neuronal dysfunction at the early stages (hypoP Tau) and late stages (hyperP Tau) of the disease.

## Materials and methods

### *C. elegans* strains and maintenance

*C. elegans* were maintained as previously described (Stiernagle, 2006). Briefly, worms were kept on Nematode Growth Medium (NGM) streaked with E. coli OP50 as food source. All assays were performed at 20°C. The N2 Bristol strain, as well as CB307(*unc-47(e307)*), DA465 (*eat-2(ad465)*), CB246 (*unc-64(e246)*) and CB1071(*unc-29(e1072)*) were obtained from the *C. elegans* Genetics Center (University of Minnesota, Minneapolis), which is funded by NIH Office of Research Infrastructure Programs (P40 OD010440). IZ629 (*ufIs34[Punc-47p::mCherry]*) was a kind gift from Dr. Michael M. Francis (University of Massachusetts, Worcester, MA). The transgenic strains were generated by Knudra Transgenics by injecting two plasmids, one with the human protein tau (0N4R) under the control of *unc-47* promoter, the second is a co-injection “marker” plasmid that expresses GFP or mCherry under the control of a *myo-2* promoter. To mimic the hypophosphorylation or the hyperphosphorylation of the tau protein, we induced a mutant carried of 12 mutations on sites close to the prolin-rich region to replace the amino-acid either by glutamate (E12) or alanine (A12), respectively. (see Plouffe *et al*. 2012) All of transgenic strains used were outcrossed six times to N2. GFP or mCherry positive progeny were selected and checked for the presence of human tau by PCR. Strains were sequenced using the following primers: 5’ GCT CTT ATC CCA CTT TGT ACA AGA AAG CT 3’ and 5’ CGC CAC CAG GAT TCC AGC AAA 3’.

### Western <u>blotting</u>

Ten plates of worms for each strain were collected in M9 buffer and protein extraction was performed in Lysis buffer (50mM HEPES pH 7.4, 1mM EGTA, 1mM MgCl2, 10% glycerol, 0.05% NP-40 and a Protease inhibitor cocktail tablets). Worms were sonicated on ice for 20sec at 30% 3 times and one more time at 40%, centrifuged at 16 000 g for 20 minutes at 4°C. The proteins were quantified using Bio-Rad DC Protein assay (Bio-Rad Laboratories). Sixty micrograms of proteins were loaded in a 10% acrylamide gel at 30mA and transferred to nitrocellulose membranes. Membranes were immunoblotted with the following primary antibodies: polyclonal rabbit Anti-Human Tau (1:10 000, Dako) and monoclonal mouse Anti-Actin (1:25 000, MP Biomedical). All the secondary antibodies used to reveal the primary antibodies were coupled with HRP, used at 1:10 000 and were purchased from Jackson ImmunoResearch. For quantification, western blot image acquisition was performed using a ChemiDoc MP system (Bio-Rad Laboratories) and densitometry analysis was done with Image Lab software (version 5.0, Bio-Rad Laboratories).

### Lifespan assays

Approximately 120 age-synchronized worms for each strain (3 plates of 40 worms each) were tested every 2 days from day 1 adult until death. Worms were scored as dead if they failed to respond to tactile stimulus and showed no spontaneous movement or response when prodded. The dead worms displaying internally hatched progeny, extruded gonads, or that crawled off the plate were excluded. All experiments were done at 20°C and each condition was done in triplicates.

### Paralysis assay on solid media

Briefly, 40 age-synchronized worms were transferred to NGM plates and scored daily for paralysis, from day 1 to day 12 of adulthood. Animals were counted as paralyzed if they failed to move upon prodding with a worm pick. Worms were scored as dead if they failed to move their head after being prodded on the nose and showed no pharyngeal pumping. All experiments were conducted at 20 °C and in triplicates, three times.

### Fluorescence microscopy

For scoring of neuronal processes for gaps or breakage, worms were selected at day 1, 5 and 9 of adulthood for visualization of motor neuron *in vivo*. Animals were immobilized in 5 mM levamisole dissolved in M9 and mounted on slides with 2% agarose pads. At least one hundred worms were scored per condition, over 4 distinct experiments.

### Progeny assay

L4 hermaphrodites were placed on individual petri dishes and transferred every day to new dishes until the death of the animal or when there was no progeny production anymore. Progeny were counted two days after transfer. 10 animals were used for each strain. Experiments were done in triplicate. All experiments were done at 20°C.

### Pumping assay

Pharyngeal activity was scored at day 1 of adulthood on 10 worms by measuring frequency of pharyngeal pumping. A pharyngeal pump was considered as one contraction-relaxation cycle of the terminal bulb of the pharyngeal muscle. The number of pharyngeal pumps was visually scored for 30 sec, over 3 trials. Experiments were done in triplicate. The mean of pharyngeal pumping rate was calculated for the 3 trials and expressed as pumping rate per minute.

### Liquid culture motility assays

Synchronized day 1 adult animals were transferred into 100 uL of M9 buffer in a 96 well plate, to a number of 30 animals per well. The 96 well plates were put into a PhylumTech WMicrotracker-One for automated data analysis over a period of 10 hours.

### *WormLab*® (worm length, swimming speed, initiation wave rate)

Worms were transferred on NGM plates with a central dot of OP50, forcing the worms to stay at the center of the plate. To induce swimming movement, 100µL of M9 was added at the center of the plates. Swimming movement was then recorded at day 1, 5, and 9 of adulthood. Animals were recorded for 30 seconds with Stereomicroscope Leica S9i with a camera CMOS integrated. Videos were taken without adding M9 to measure worm length. For the analysis, the WormLab® software was used to calculate the swimming speed, initiation wave rate and the length of each worm. All experiments were conducted at 20°C. Experiments were done in triplicate.

### Aldicarb sensitivity assay

Worms were grown on standard NGM plates until day 1 of adulthood, when they were transferred to NGM plates containing 1 mM aldicarb. Worm paralysis was assayed every 30 minutes for 2 hours. Animals were counted as paralyzed if they failed to move upon prodding with a platinum wire. Around 300 worms were tested for each condition, over 3 trials.

### Levamisole assay

Worms were grown on standard NGM plates until day 1 of adulthood, when they were transferred to NGM plates containing 0.25 mM levamisole. Worm paralysis was assayed every 30 minutes for 2 hours. Animals were counted as paralyzed if they failed to move upon prodding with a platinum wire. Approximately 300 worms were tested for each condition, over 3 trials.

### Statistics and reproducibility

All experiments were repeated at least three times. Quantitative data were expressed as mean ± SEM. GraphPad Prism v8 software was used for all statistical analyses. All statistic tests and experimental n are clearly indicated in figure legends.

## Acknowledgements

This work was supported by Canadian Institute of Health Research (CIHR) grants to N.L. and J.A.P.. We thank all the laboratory members for helpful discussions.

## Author contributions

A.L., E.S., J.P., S.B., M.L. and C.M. performed the experiments and analyzed the data. A.L., N.L. and J.A.P. contributed to manuscript editing.

## Competing interest statement

The authors declare no competing interests.

## Notes

### Competing Interest Statement

The authors have declared no competing interest.

