## Supplementary figures for "Mimicking of tau hyperphosphorylation in GABAergic motoneurons of *C. elegans* induces severe peripheral and neuronal alterations"

**Figures S1. Plasmid constructs for Tau expression in *C. elegans*.** Expression vector used for GABAergic expression of human Tau under the *unc-47* promotor. The following full-length Tau constructs were cloned and injected into *C. elegans*: **(A)** WT Tau, **(B)** HypoP Tau containing 12 mutated sites to alanine, and **(C)** HyperP Tau containing 12 mutated sites to glutamate. Restriction sites are indicated on the plasmids.

**Figures S2. Confirmation of mutations in Tau sequence. (A)** Sequence alignment of WT Tau (identified seq_1, 0N4R Tau) and HypoP Tau (identified seq_2). # represents mutations in original sequence, leading to change in amino acids (alanine, A), indicated by green arrows. **(B)** Sequence alignment of WT Tau (identified seq_1, 0N4R Tau) and HyperP Tau (identified seq_2). # represents mutations in original sequence, leading to change in amino acids (glutamate, E), indicated by red arrows.

**Figures S3. Tau transgenics do not display any pharyngal pumping impairment.** *eat-2*(ad465) mutant was used as positive control for pharyngal pumping deficits. Neither WT, HypoP nor HyperP Tau transgenics showed differences in pumping rate (per minute). One-way ANOVA was performed. 30 individual worms were tested for each condition. For boxplots, minimum, first quartile, median, third quartile, and maximum are shown. ****p < 0.0001.

**Figures S4. HyperP Tau induces decreased initiation wave rate.** Wave initiation rate of nematodes at day 1 **(A)**, 5 **(B)** and 9 **(C)** of adulthood. Wave initiation rate is the number of body waves initiated by the head or the tail per minute of swimming. HyperP Tau wave initiation rate is significantly lower compared to N2 worms at day 1, 5 and 9 of adulthood.

HypoP worms have an higher wave initiation rate at day 1, but not at day 5 or 9. One-way ANOVA was performed. Between 88 and 188 individual worms were tested for each condition. For boxplots, minimum, first quartile, median, third quartile, and maximum are shown. ***$p < 0.001$; ****$p < 0.0001$

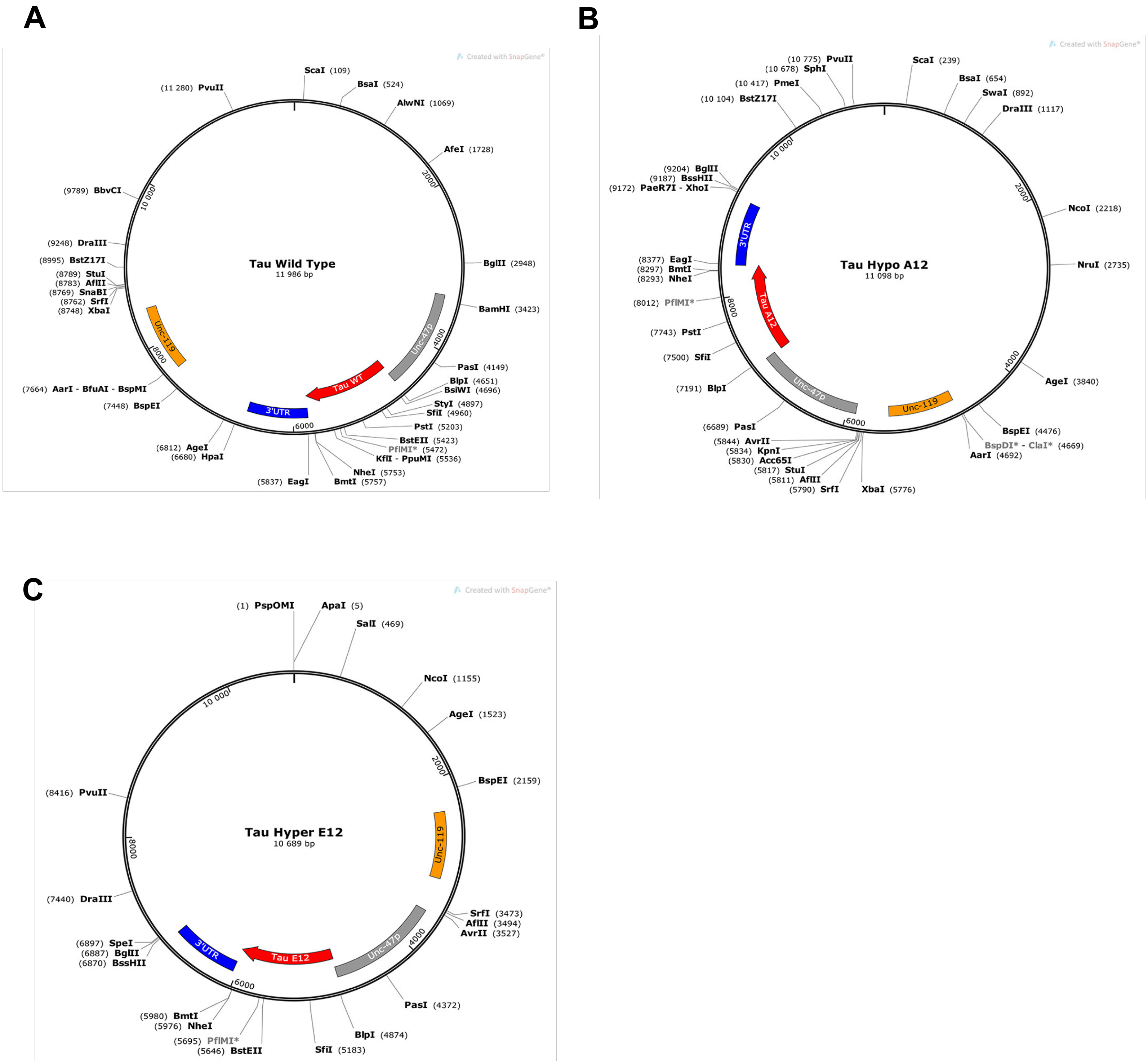
A
Tau Wild Type
11 986 bp
ScaI (109)
BsaI (524)
AlwNI (1069)
AfeI (1728)
BglII (2948)
BamHI (3423)
PasI (4149)
BlpI (4651)
BsiWI (4696)
StyI (4897)
SfiI (4960)
PstI (5203)
BstEII (5423)
PflMI* (5472)
KflI - PpuMI (5536)
NheI (5753)
BmtI (5757)
(5837) EagI
(6812) AgeI
(6680) HpaI
(7448) BspEI
(7664) AarI - BfuAI - BspMI
(8748) XbaI
(8762) SrfI
(8769) SnaBI
(8783) AflII
(8789) StuI
(8995) BstZ17I
(9248) DraIII
(9789) BbvCI
(11 280) PvuII
Unc-119
Unc-47p
Tau WT
3'UTR
2000
4000
6000
8000
10 000
Created with SnapGene®
B
Tau Hypo A12
11 098 bp
(10 775) PvuII
(10 678) SphI
(10 417) PmeI
(10 104) BstZ17I
ScaI (239)
BsaI (654)
SwaI (892)
DraIII (1117)
NcoI (2218)
NruI (2735)
AgeI (3840)
BspEI (4476)
BspDI* - ClaI* (4669)
AarI (4692)
XbaI (5776)
(5790) SrfI
(5811) AflII
(5817) StuI
(5830) Acc65I
(5834) KpnI
(5844) AvrII
(6689) PasI
(7191) BlpI
(7500) SfiI
(7743) PstI
(8012) PflMI*
(8293) NheI
(8297) BmtI
(8377) EagI
(9172) PaeR7I - XhoI
(9187) BssHII
(9204) BglII
3'UTR
Tau A12
Unc-47p
Unc-119
2000
4000
6000
8000
10 000
Created with SnapGene®
C
Tau Hyper E12
10 689 bp
(1) PspOMI
ApaI (5)
SalI (469)
NcoI (1155)
AgeI (1523)
BspEI (2159)
SrfI (3473)
AflII (3494)
AvrII (3527)
PasI (4372)
BlpI (4874)
SfiI (5183)
(5646) BstEII
(5695) PflMI*
(5976) NheI
(5980) BmtI
(6870) BssHII
(6887) BglII
(6897) SpeI
(7440) DraIII
(8416) PvuII
Unc-119
Unc-47p
Tau E12
3'UTR
2000
4000
6000
8000
10 000
Created with SnapGene®

**Figure S1**

A

WT Tau vs HypoP Tau

B

WT tau vs HyperP Tau

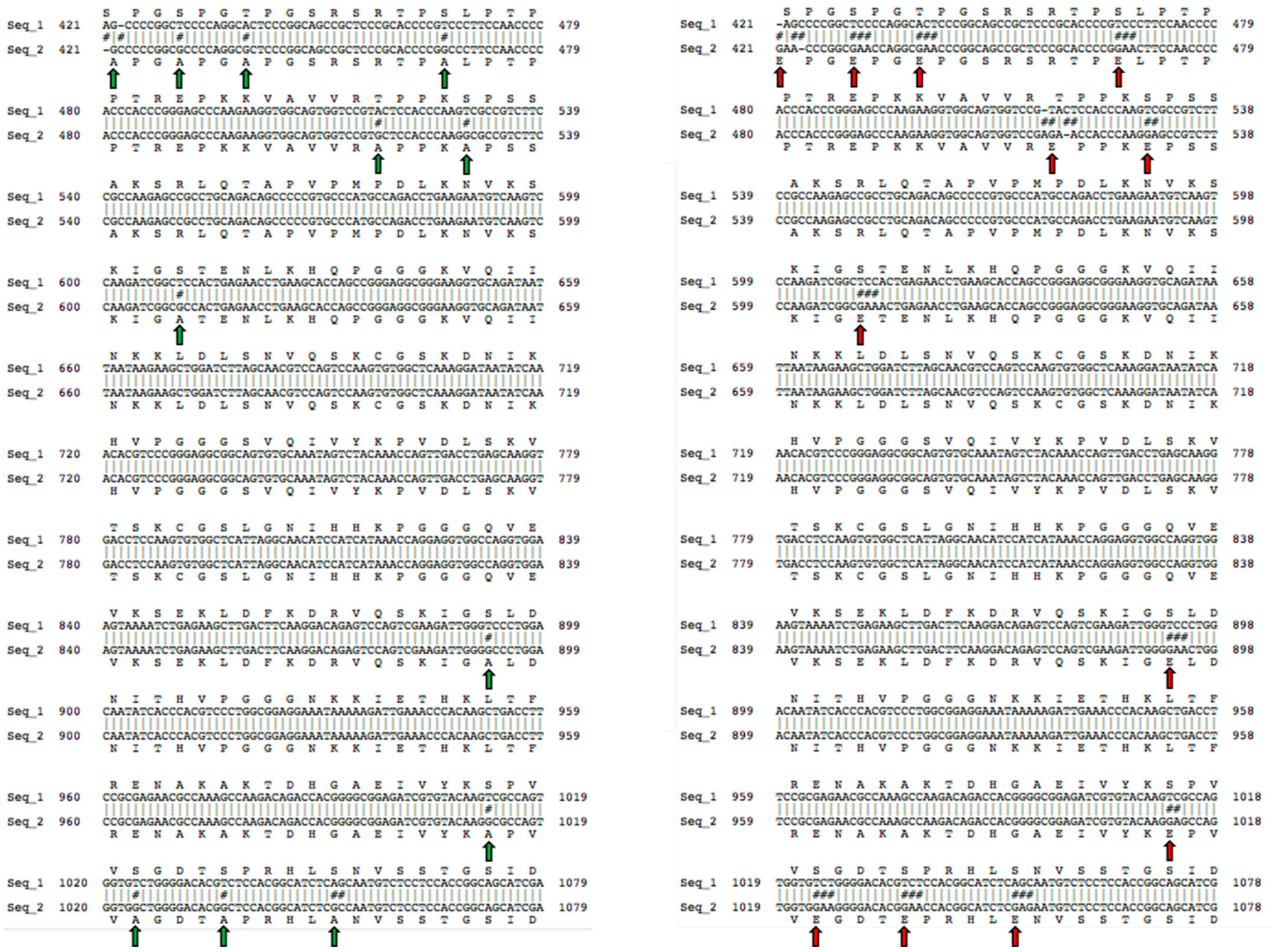

**Figure S2**

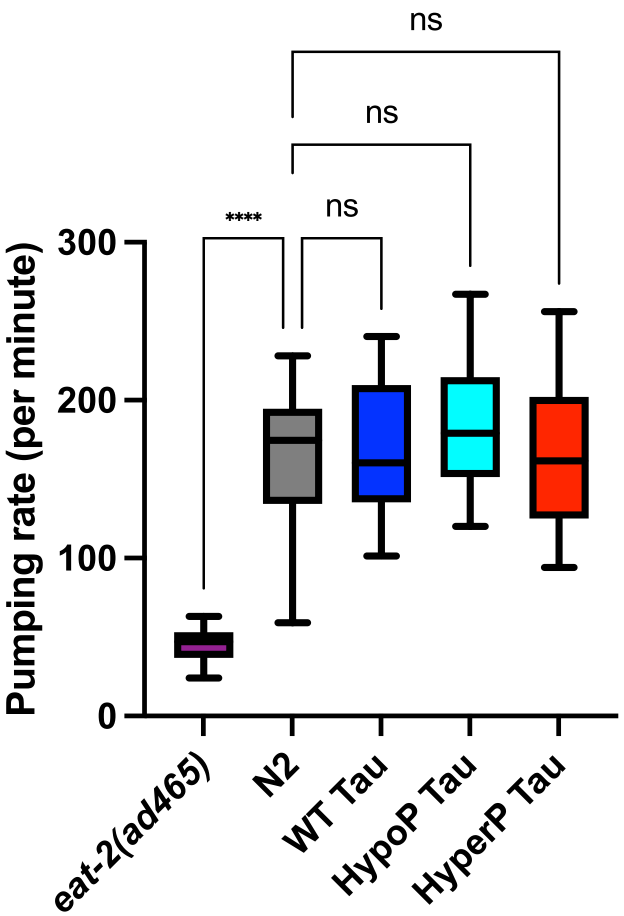
ns
ns
****
ns
Pumping rate (per minute)
300
200
100
0
eat-2(ad465)
N2
WT Tau
HypoP Tau
HyperP Tau

Figure S3

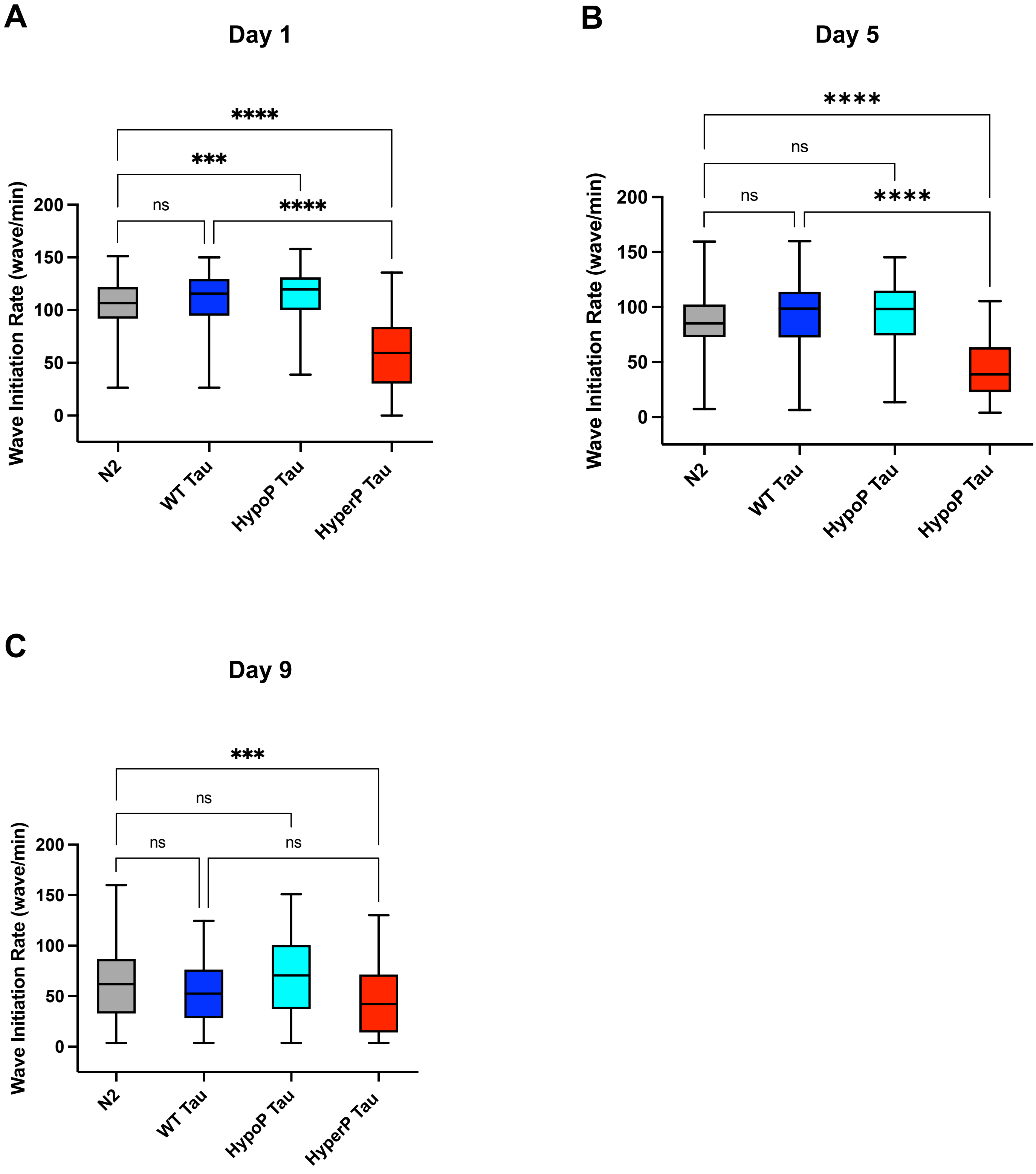

A
Day 1
****
***
ns
****
Wave Initiation Rate (wave/min)
200
150
100
50
0
N2
WT Tau
HypoP Tau
HyperP Tau
B
Day 5
****
ns
ns
****
Wave Initiation Rate (wave/min)
200
150
100
50
0
N2
WT Tau
HypoP Tau
HypoP Tau
C
Day 9
***
ns
ns
ns
Wave Initiation Rate (wave/min)
200
150
100
50
0
N2
WT Tau
HypoP Tau
HyperP Tau


**Figure S4**
